# Investigating antiviral pathways in Atlantic salmon cells through interferon receptor knockouts via CRISPR-Cas9

**DOI:** 10.1101/2025.01.14.632998

**Authors:** Mohammad Ali Noman Reza, Thomas Nelson Harvey, Amr Ahmed Abdelrahim Gamil, Øystein Evensen, Gareth Benjamin Gillard, Guro Katrine Sandvik

## Abstract

In Atlantic salmon (*Salmo salar*), infectious salmon anemia virus (ISAV) and infectious pancreatic necrosis virus (IPNV) evade host immune response through complex antagonistic mechanisms. Type I interferons (IFNs) play a pivotal role in antiviral defense by signaling through heterodimeric receptors to activate the JAK-STAT pathway and drives the expression of interferon-stimulated genes (ISGs). In this study, CRISPR-Cas9 was used to knock out (KO) interferon receptor genes (*crfb1a*, *crfb5a*, *il10rb*, *ifngr2a*) and a combined group of candidate receptors (*crfb1a*, *crfb5a*, *il10rb*, *ifngr2a*, *il10r2*) to investigate their roles and their impact on downstream signaling cascades with RNA sequencing. Recombinant IFNa was used to induce an antiviral state before challenging cells with ISAV and IPNV.

The knockouts significantly disrupt downstream antiviral signaling, with two knockouts, *crfb1a* and *crfb5a*, showing pronounced effects. During ISAV infection, the *crfb1a* KO group exhibited a marked reduction in the expression of critical signaling genes such as *stat1b, stat2, stat6,* and *irf3* during ISAV infection, while *irf7* was upregulated during IPNV infection. The *crfb5a* KO group exhibited reduced *stat2* expression in ISAV infection and upregulated *irf7* during IPNV infection. Despite these disruptions, ISGs such as *Mx* and *isg15* maintained their expression levels across all knockout groups, suggesting potential alternative signaling pathways. Pathway analysis further revealed upregulation of cellular processes like actin regulation and phagosome activity, which may compensate for impaired immune signaling. These findings highlight the distinct roles of IFN receptor genes in mediating antiviral responses and underscore the complexity of IFN signaling in Atlantic salmon.

## Introduction

Atlantic salmon (*Salmo salar*) is the most extensively farmed cold-water fish species globally (FAO, 2024) and the aquaculture production has reached approximately 2.87 million metric tons in 2023. However, high densities of fish facilitate disease spread and infectious diseases has been among the main challenges facing the industry. Vaccination has helped in controlling many of the bacterial diseases but most of the viral diseases such as infectious salmon anemia virus (ISAV) and infectious pancreatic necrosis virus (IPNV) has remained as a persistent challenge.

Virus infect host cells by interacting with the host cell molecules, or receptors (Thorley et al., 2010). Once inside the cells, viral RNA released into the cytoplasm may be transported into the nucleus where the transcription and replication occurs as observed in ISAV (Eliassen et al., 2000). For other viruses such as IPNV, however, the transcription and replication occurs in cytoplasm (Skjesol, 2009). During the intracellular replication phase, viral RNA can be detected by pattern recognition receptors (PRRs) such as transmembrane Toll-like receptors (TLRs) or cytosolic retinoic acid-inducible gene I (RIG-I)-like receptors (RLRs) (Loo & Gale, 2011; Svingerud et al., 2013). These PRRs recognize pathogen-associated molecular patterns (PAMPs) and trigger signaling cascades that lead to the transcription of type I interferon genes, such as IFNa and IFNb, IFNc.

Interferons (IFN) are cytokines which are essential defense mechanism in fish against virus infections (Robertsen, 2006). These cytokines exhibit their effects by binding to unique IFN receptors on the immune cell surface, leading to the induction or increase expression of interferon stimulated genes (ISGs). Once IFNa binds to its receptor, the associated JAKs (JAK1, Jak2/Tyk2) come into close proximity and activate each other through cross-phosphorylation (Dehler et al., 2019; Gan et al., 2020; Shemesh et al., 2021). The phosphorylated JAKs then phosphorylate STAT1, which forms homodimers or heterodimers with STAT2 (Gan et al., 2020). This dimer, together with IRF9, forms the interferon-stimulated gene factor 3 (ISGF3) complex which translocate to the nucleus. In the nucleus it binds to Interferon-stimulated response elements (ISREs) and induces ISGs expression. It binds with the interferon-stimulated response elements (ISREs) and induce the ISGs expression. These ISGs, including *Mx*, *isg15*, and *pkr*, are essential antiviral proteins that inhibit virus replication and spread to neighboring cells (Varela et al., 2014).

While three types of IFNs (I-III) are identified in mammals, only type I and type II of IFNs have been identified in fish (Martin et al., 2007). Salmonid type I interferons consists of large numbers of related proteins including IFNa, IFNb and IFNc, which specifically works against virus (Takaoka & Yanai, 2006; Zou et al., 2007) and type II consists of a single protein IFN gamma (IFN-γ), primarily known for its role in activating macrophages (Young & Hardy, 1995).

In general, IFNs exert their functions by binding to heterodimeric IFN receptors on the cell surface, activating the JAK-STAT signaling pathway and inducing the expression of ISGs. The IFN receptors in fish classified as class II cytokine receptor family member (CRFB) either with a long or a short cytoplasmic chain (Lutfalla et al., 2003). In CHSE-24 cells, Atlantic salmon IFNa can signal through binding to long chain CRFB1a, with short chain CRFB5a, CRFB5b, or CRFB5c (Sun et al., 2014). The long chain CRFB1a has been suggested to play a crucial role in forming functional receptors for IFNs while CRFB5a, CRFB5b, CRFB5c short chains are thought to help in regulating host antiviral and antimicrobial functions (Sun et al., 2014; Wang et al., 2023).

In this study, we used CRISPR-Cas9 to knock out (KO) multiple Atlantic salmon candidate IFN receptor genes *crfb1a*, *crfb5a*, *il10rb*, *ifngr2a*, and a combined group of candidate receptors (*crfb1a*, *crfb5a*, *il10rb*, *ifngr2a*, *il10r2*) in SHK-1 cells. Subsequently, the cells were treated with recombinant IFNa both in KO and no_KO groups to keep the cells in an antiviral state before challenging them with ISAV and IPNV. Our aim was to determine how IFN receptor KOs affect downstream pathways, IRF signaling, and ISG production.

## Material and methods

### Cell culture, virus propagation and titration

The cell culture, virus propagation and virus titration for this study was based on the method presented in Reza et al., 2024. The SHK-1 (ECACC 97111106) and ASK (ATCC CRL-2747) cells were obtained from Øystein Evensen of the Norwegian University of Life Sciences. For culturing cells, culture media was composed of Leibovitz L-15 media with Glutamax, 10% Fetal Bovine Serum (FBS: non-USA origin, sterile-filtered, Sigma-Aldrich) and 1% pen-strep (Penicillin-Streptomycin, Thermofisher). When reaching 80-90% confluency, the cells were detached using 0.05% trypsin/EDTA, subcultured and then maintained at 20°C under atmospheric conditions.

For virus propagation ASK cells were seeded in 25 cm^2^ flasks and grown until 70%-80% confluenc. Cells were then inoculated with ISAV and virulent IPNV (T_217_A_221_) isolate. During the virus infection, the culture medium contained 1% FBS and 1% pen-strep throughout the infection period. Cells were incubated for 7-14 days at 15°C until cytopathic effect became apparent. The culture media was collected, filtered with 0.20 µM filter and allocated in 2 ml cryopreservation tube and stored in Cryofreezer (-150 °C). Virus titer was determined using the 50% tissue culture infective dose (TCID50), following the procedure described by Kärber, 1931.

SHK-1 cells which has macrophage-like characteristics (Dettleff et al., 2024), were used for immunological studies and ASK cells which, are epithelial type cells (Ørpetveit et al., 2008). were used for the propagation of ISAV and IPNV.

### IFN receptor selection

In Atlantic salmon type I IFN receptors (CRFBs) are present in two cluster in salmon chromosome 21 and chromosome 25, where the first cluster has *crfb1a*, *crfb3*, *crfb4a* (*il10r2a*), *crfb5c*, *crfb5x* and a *ifngr2* related gene and the second cluster has *crfb5a* and an *ifngr2* gene (Sun et al., 2014). In this study, we have used RNP (ribonucleoprotein) mediated CRISPR- Cas9 method to create KO cell lines *crfb1a* (LOC106581740), *crfb5a* (LOC106581895), *il10rb* (LOC106581741), *ifngr2a* (LOC106581880), and a combined group of receptors (*crfb1a*, *crfb5a*, *il10rb*, *ifngr2a, il10r2* [LOC106586857]) from the SHK-1 cell line.

### Producing IFN receptors KO cell lines

Three crRNAs were designed in silico for each genes utilizing the Synthego KO guide design tool (https://design.synthego.com/). We used this tool to design crRNAs that target early exons of the protein-coding transcript, which are shared across most transcripts and located within the coding sequence. However, in the case of *crfb5a*, two transcript variants begin in very late exons. For this gene, we designed gRNAs targeting the 3rd exon, even though one transcript begins at the 7th exon, assuming that targeting the earlier exon would not significantly impact functionality. We also ensured that each gRNAs had the high predicted activity scores defined by Doench et al. (2016) and no potential off-target sites with 0 or 1 base mismatches. The crRNAs and tracrRNA were ordered from Integrated DNA Technologies (IDT).

RNP complexes were assembled by modifying Gratacap et al., 2020 protocol to transfect the gRNA and Cas9 into the cells. Briefly, obtained crRNAs and tracrRNA were resuspended in nuclease free water to get 100 µM. Equal volumes of crRNA and tracrRNA were mixed in a tube to prepare gRNA. The tube was incubated at 95° C for 5 minutes and cooled on the benchtop. Next, we added 20 µM of Cas9 (NEB, Ipswich, USA) in the same volume as the gRNA to form the RNP complex (Ribonucleoprotein complex). So, the final concentration became 10 μM of Cas9 and 25 μM of gRNA. The RNP complex was then incubated at room temperature for 15 minutes and transferred to ice until further use. For electroporation, a mixture of 1.4 µl RNP and 2.6 µl of OptiMEM reduced serum media (Gibco) was prepared.

Next, SHK-1 cells were prepared for the electroporation. For this, confluent flask of SHK-1 cells were sub cultured and grown in a new flask with fresh media. In the third day after subculture, cells were harvested using trypsinization, counted, and centrifuged for 5 minutes at 200g. The supernatant was removed carefully, and the cells were washed by resuspending in 2 ml of PBS, followed by a second centrifugation at 200g for 5 minutes. The PBS was then removed, and cells were resuspended in OptiMEM reduced serum media to a final concentration of 1*10^7 cells/ml. For each electroporation, 10 µl of this cell suspension was added to the RNP mixture, resulting in a final concentration of cas9 1 µM and gRNA 5 µM and incubated at room temperature for 5 minutes. The cell-RNP mixtures were then electroporated at 1400 V, 20 ms, and 2 pulses using the Neon Transfection System and 10 µl Kit (Invitrogen, USA). The electroporated cells were seeded into 24-well plates with fresh culture medium without antibiotics. The following day, the medium was replaced with one containing 1% pen strep. We also attempted to produce cells with all five receptors knocked out simultaneously. For this we took 1.4 µl of RNP for each receptor, and 13 µl of OptiMEM reduced serum media (2.6 µl per receptor) was added to the combined mixture. The solution was thoroughly mixed, and 4 µl of the RNP mixture was added to 10 µl of cells. A total of 10 µl of this final mixture was used for the transfection.

### Assessing CRISPR-Cas genome editing efficiency and specificity via Sanger sequencing

To assess the KO efficiency of each gRNA, cells were harvested seven days post-electroporation. The cells were detached using 100 µL of trypsin, followed by the addition of 400 µL of medium to neutralize the trypsin. A 100 µL aliquot of the cell suspension was then taken for genomic DNA extraction, while the remaining 400 µL of cells were recultured. For DNA extraction we used QuickExtract™ DNA Extraction Solution (Lucigen, Middleton, USA) by making modification of the manufacturer’s protocol. Briefly, 30 μL of QuickExtract buffer was added to the cells, and mixed thoroughly by pipetting, and then incubated at room temperature for 5 minutes. This was followed by incubation at 65 °C for 15 minutes and 98 °C for 2 minutes using a PCR machine.

To amplify the target site for sequencing, a single primer pair was designed to produce a product covering the region containing the three gRNAs tested for each gene (Table 1). PCR amplification was performed in 50 μL reaction volumes using Q5® High-Fidelity 2X Master Mix (NEB) and 1 μL of genomic DNA. The thermal cycling conditions included an initial denaturation at 98°C for 30 seconds, followed by 37 cycles of denaturation at 98°C for 10 seconds, annealing at 60°C for 15 seconds, and extension at 72°C for 15 seconds per kilobase. A final extension step was carried out at 72°C for 2 minutes, with a final hold at 4°C. To verify the amplification, a portion of the PCR product was electrophoresed on a 1% agarose gel to confirm the presence of a single band of the correct size. Upon confirmation, PCR purification was done by using, ExoSAP-IT (Applied Biosystems™) by following manufacturers protocol with modifications. The reagent was diluted to 1:200 and 1 µl of the diluted reagent was used for purification of 5 µl pcr product. Subsequently, 5 µM of sequencing primer was added to 5 µl of the purified PCR product, and the mixture was submitted for Sanger sequencing by using Mix2Seq (Eurofins Genomics). Obtained chromatogram result (.abi files) was analyzed by using the Inference of CRISPR Edits (ICE, Synthego Inc) software. Best gRNA for each receptor was selected based on their indel efficiency.

**Table 1:**
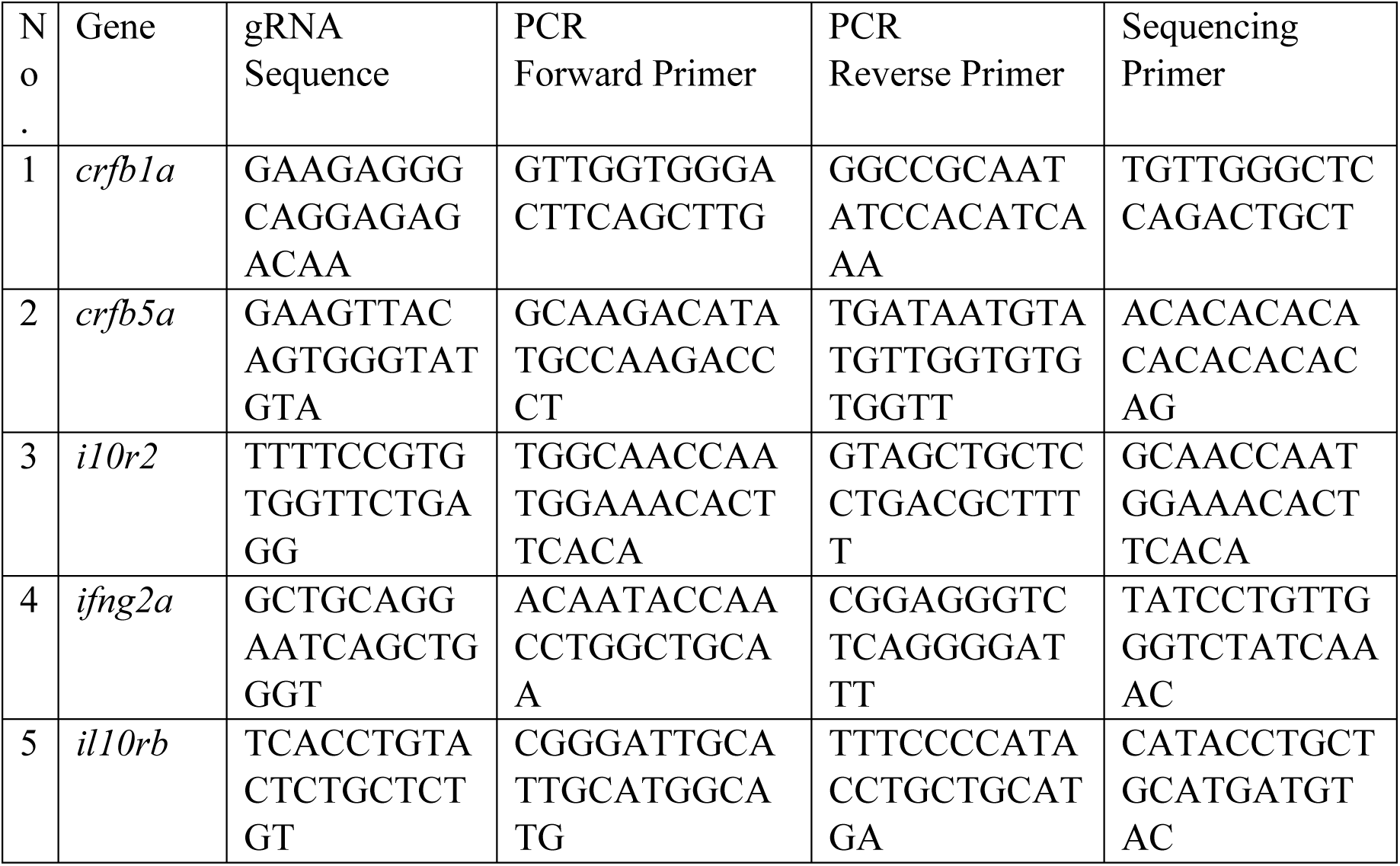
PCR Primers and Sequencing Primers for CRISPR-Cas9 Targeted Genes.

Cells lines with higher KO percentage were maintained and transferred to T-25 flasks once confluent. We then proceeded to the IFNa experiments, confirming KO proportion before each experiment as described above to ensure the KO population remained stable.

### IFNa treatment

Each group of KO cells along with wildtype SHK-1 cells were plated in 48-well plates at a density of 2×10^5 cells. The following day after cell adherence the media was removed, cells were washed with phosphate-buffered saline (PBS) and fresh media with 1% fetal bovine serum (FBS) was added. Two experimental groups were formed, one with IFNa treatment (235ng/ml) and one without (no IFNa).

After 24 hours, KO and WT cells were further divided into three subgroups, one uninfected control, one challenged with infectious salmon anemia virus (ISAV) at a multiplicity of infection (MOI) of 0.20, and one challenged with infectious pancreatic necrosis virus (IPNV) at an MOI of 0.10. In all cases, infection (or mock infection) lasted for 2 hours. The cells were then washed with PBS and collected for RNA extraction. For this, 100 µL of TRI reagent (Direct-zol RNA Microprep Kit, Zymo) was added to the cells, which were carefully scraped using a scraper and collected into a tube. The tube was immediately frozen at -80 °C.

### RNA extraction

Briefly, total RNA was extracted by using by using Direct-zol RNA Microprep Kits (Zymo), following manufacturer’s protocol. The purity and the concentration were assessed by using NanoDrop 8000 Spectrophotometer (Thermofisher Scientific) and NanoDrop 8000 V2.1.0 software and Agilent 4150 TapeStation system (Agilent) was used to check the RNA integrity. The sample was considered pure if the ratio is around 2.0 and standard quality where RIN is greater than 8.

### RNA-seq and data analysis

To understand the impact of knocking out the different interferon alpha (IFNa) receptors in more detail, RNA sequencing analysis was conducted on IFNa-induced ISAV or IPNV-infected SHK-1 cells with various receptor KOs (n=3). The results were compared with those from IFNa_ISAV/IFNa_IPNV treatments in non-KO SHK-1 cells (n=4). Briefly, library preparation and sequencing were performed by Novogene Co., LTD (UK), using a stranded mRNA library preparation method. The sequencing was executed on an Illumina platform (Illumina NovaSeq 6000), which generate paired end (PE)150 bp reads. Each sample was sequenced to a depth of around 20 million reads which ensures the coverage for accurate transcript quantification and The raw reads in FASTQ format were processed using the nf-core/rnaseq pipeline (Ewels et al., 2020) which includes Trim Galore, FastQC/MultiQC, STAR aligner, Salmon and FeatureCounts. We used Atlantic salmon reference genome Ssal_v3.1 for the alignment. The RNA-seq data was analyzed using custom scripts and configurations run on the Orion HPC clusters.

## Result

### Temporal KO efficiency of IFN receptor genes in SHK-1 cells using single and combined gRNAs

KO efficiency of different gRNAs for *crfb1a*, *crfb5a*, *il10rb*, *ifngr2a*, and *il10r2* in SHK-1 cells were measured using insertion/deletion (indel) percentages at multiple time points post-electroporation (days 7, 14, 21, 92, and 104). Discordance plot and indel frequency distribution plot showed KO efficiency for different IFN receptors across different time points (Figure 1).

**Figure 1:**
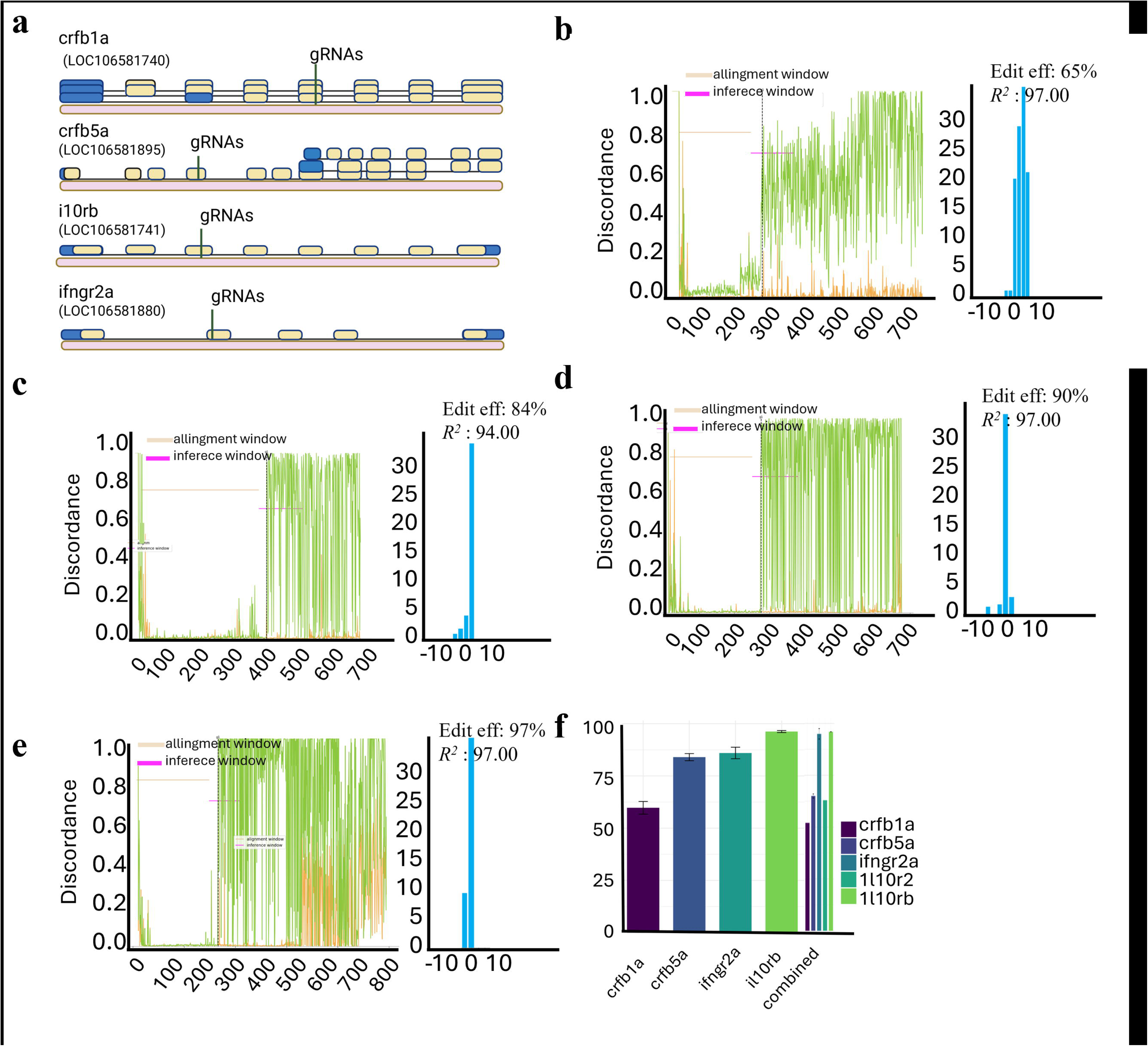
Designing guides for CRISPR Cas9 KO of IFN receptors in SHK-1 cells. 1.a Position of gRNAs (BioRender) 1b-e. Discordance plot and indel frequency distribution plot for best gRNA of each receptor of a specific date. 1f Bar graph illustrating the average knocKOut efficiency of each gRNA at different time points post-transfection, as measured by indel percentage. Error bars represent standard deviation (SD).

In *crfb1a*, the gRNA showed moderate editing efficiency of 65% after 7 days of transfection, however gradually declined over time and showing 51% knockout cells after 92 days. (Figure 1b, S.T1). C*rfb5a* gRNA showed 78% editing efficiency after 7 days becoming 88% and 86% after 14 and 104 days respectively (Figure 1c, S.T1). The gRNA for *ifngr2a* produced consistently high KO efficiency (90%) from 7 to 104 days post transfection (Figure 1d, S.T1). Similarly, *il10rb* showed higher editing efficiency (90%) over different time (Figure 1e, S.T1). Simultaneous KO of the IFN receptor genes (*crfb1a*, *crfb5a*, *il10rb*, *ifngr2a*, and *il10r2*) was also evaluated over time (Figure 1f, S.T2). At 7 days post-transfection, *crfb1a* showed a modest KO efficiency of 52%, while *crfb5a* and *ifngr2a* demonstrated higher efficiencies of 64% and 97%, respectively. *il10rb* exhibited an efficiency of 96%, while *il10r2* reached 63%. After 31 days and 38 days of transfection the editing efficiency of different gRNAs remained similar to the one detected at 7 days after transfection. The ICE analysis also reported the frameshift mutation on each replication showing 1%-4% lower than the KO efficiency. Although most cells were edited in this experiment, a small percentage remained unedited, producing functional protein. Nonetheless, we have used the term KO of different receptors in this study to this mixed population cells.

### Transcriptomic response to IFN receptor KO

To investigate the effects of IFN receptor KO, we conducted a transcriptomic analysis using RNA-seq. Principal component analysis (PCA) (Figures 2b, c) revealed clear separation between KO (*crfb1a*, *crfb5a*, and combined KO) and no_KO groups in IFNa-induced, virus-infected samples. Since *il10rb* showed minimal clustering, and *ifngr2a* displayed inconsistent patterns, we exclude them from further analysis (S F1). Similarly, a single KO of *il10r2* was not included due to lower knockout efficiency of its guide RNA.

**Figure 2:**
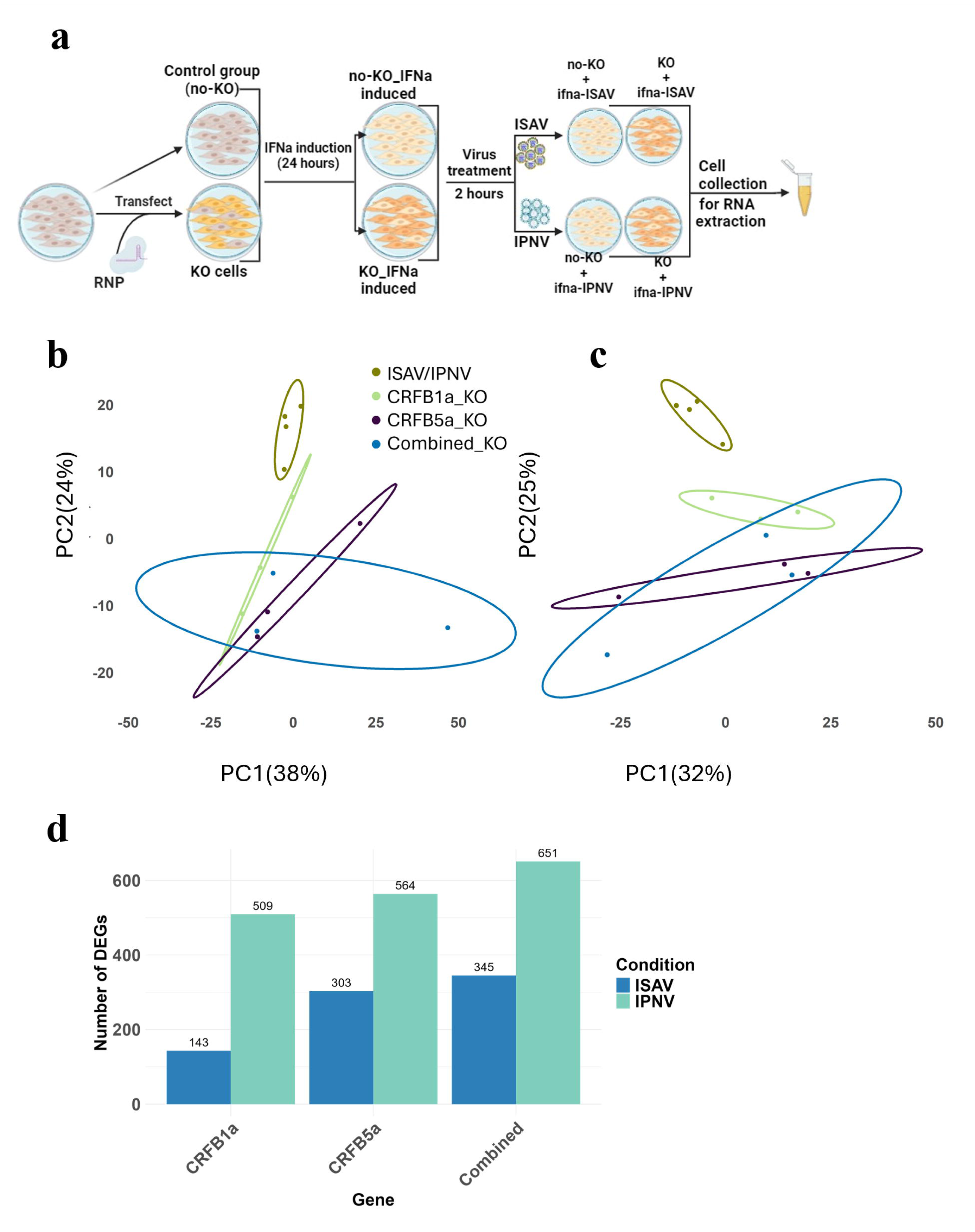
RNA Sequencing and transcriptomic analysis of SHK-1 Cells KO for IFNa receptors under ISAV and IPNV infections. 1a. Schematic representation of the experimental design, showing SHK-1 cells treated with IFNa for 24 hours followed by infection with ISAV or IPNV. SHK-1 cells were grouped into three categories: no-KO, IFNa-induced-infected-no-KO, and IFNa-induced-infected-KO. 1b,c. Principal component analysis (PCA) displaying gene expression clustering among SHK-1 cells in ISAV and IPNV cells with individual KO of crfb1a, crfb5a, and combined IFN receptor genes post-infection. PC1 and PC2 represent the indicated percentage of the variance, respectively. 1d. Bar graph illustrating the number of differentially expressed genes (DEGs) identified in response to ISAV and IPNV infections for *Crfb1a, crfb5a* and combined knockouts. The green bars represent ISAV infection, and the blue bars represent IPNV infection. Numbers on top of the bars indicate the total number of DEGs.

Differential expression analysis identified 143 DEGs in the *crfb1a* KO group infected with ISAV and 509 DEGs in the same group infected with IPNV. The *crfb5a* KO group showed 303 DEGs for ISAV and 564 DEGs for IPNV, while the combined KO group had 345 DEGs in ISAV and 651 DEGs in IPNV (Figure 2d). UpSet plots, Venn diagrams, and Volcano plots were used to identify important genes. Downregulation of genes such as *dll4, cd74a*, and *ghrb* was commonly observed across different KO groups (S F2).

### Differential gene expression in the JAK-STAT pathway

The immediate downstream pathway of IFN receptor signaling is the JAK-STAT signaling pathway. The impact of IFNa receptor KO on genes of this pathways was different in ISAV and IPNV-infected cells (Figure 3). Our study showed that in ISAV-infected cells, *crfb1a* KO led to significant downregulation of *stat1b* (*p*=0.01), *stat2* (*p*=0.003), and *stat6* (*p*=0.02). *Crfb5a* KO, on the other hand, resulted in significant downregulation of *stat2* (*p*=0.04) only. Additionally, *stat1a* showed a trend of downregulation in *crfb1a* (*p*=0.15), *crfb5a* (*p*=0.22), and combined KO (*p*=0.10) (Figure S F3) groups following ISAV infection, though these changes were not statistically significant. In IPNV-infected cells, *stat2* exhibited a downregulation trend in *crfb1a* (*p*=0.12) and *crfb5a* KO groups (*p*=0.15) (Figure 3). Another key observation was that upstream components such as *jak1* and *tyk2* showed no significant differences between the with or without KO groups in both virus-infected cells.

**Figure 3:**
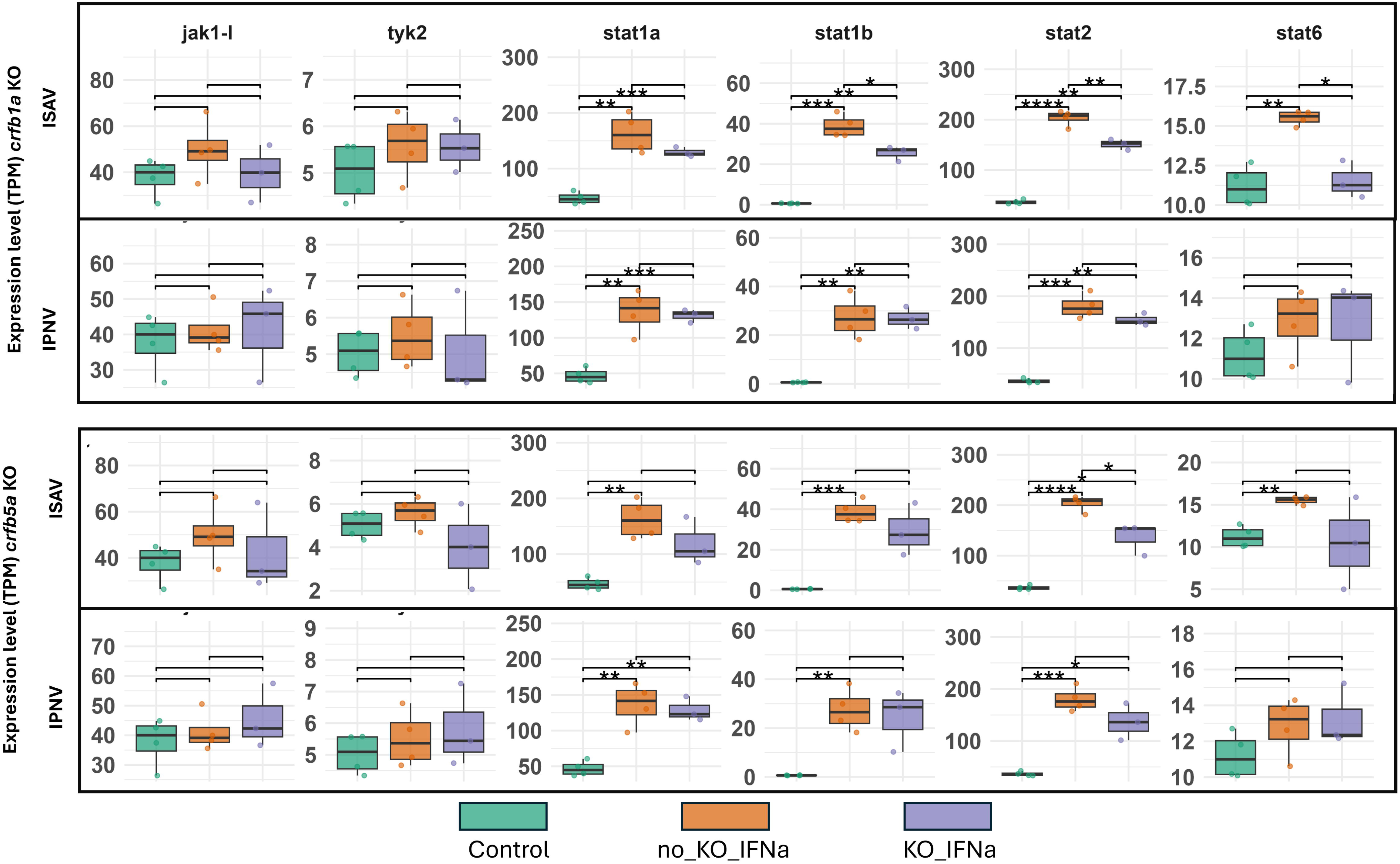
Expression levels (TPM) of key genes (*jak1-l, tyk2, stat1a, stat1b, stat2, and stat6*) in SHK-1 cells after IFNa treatment and infection. The panels represent the following conditions: (1) crfb1a knockout (KO) cells infected with ISAV (top panel), (2) crfb1a KO cells infected with IPNV (second panel), (3) crfb5a KO cells infected with ISAV (third panel), and (4) crfb5a KO cells infected with IPNV (bottom panel). Data are shown for the control group (green, non-KO without IFNa), no-KO_IFNa group (blue, non-KO with IFNa), and KO_IFNa group (orange, KO with IFNa). Statistical significance is indicated as *P < 0.05, **P < 0.01, ***P < 0.001. Error bars represent standard deviation (SD). Biological replicates: n=4 for control, n=3 for KO groups.

### Differential gene expression of genes involved in the signaling cascade

The expression of some of the essential signaling genes significantly differed between the IFNa receptor KO and no_KO groups. During ISAV infection, the transcription factor *irf3* was significantly downregulated in the *crfb1a* KO group (*p*=0.03), while its expression remained largely unaffected in IPNV-infected cells (Figure 4). In contrast, *irf7* displayed an upward trend in expression in *crfb1a* (*p*=0.14) and *crfb5a* (*p*=0.06) KO groups during ISAV infection compared to the non-knockout group, although these changes were not statistically significant. In IPNV-infected samples, however, *irf7* was significantly upregulated in both the *crfb1a* (*p*=0.01) and *crfb5a* (*p*=0.03) KO groups (Figure 4). Interestingly, *irf1_isoform2_*, which showed no significant change in ISAV-infected cells, was significantly reduced in *crfb1a* KO samples (*p*=0.02) and also showed a downregulation trend in *crfb5a* KO cells (*p*=0.07) during IPNV infection, though the latter trend was not statistically significant.

**Figure 4:**
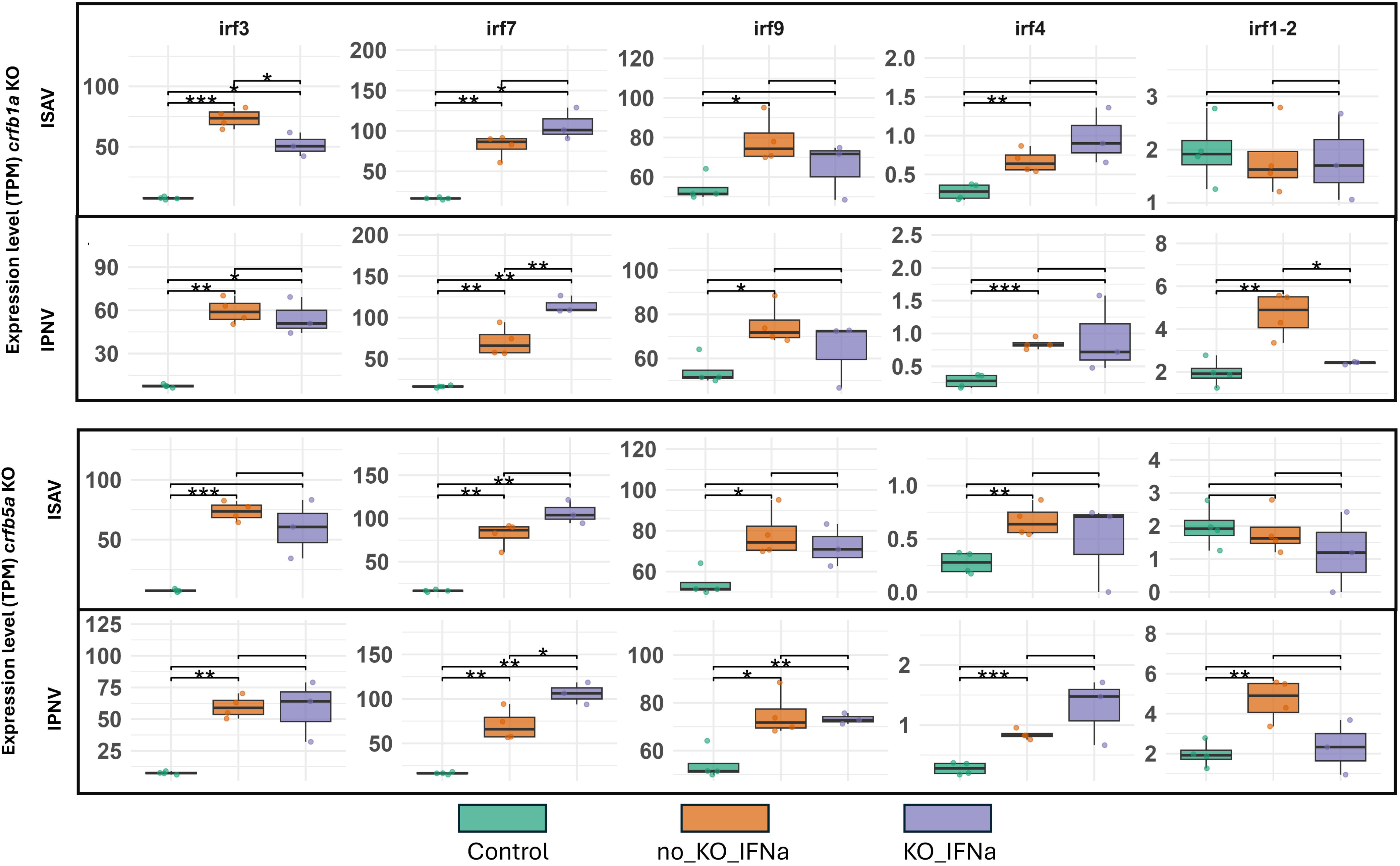
Expression levels (TPM) of key involved in the signaling cascade (*irf3, irf7, irf9, irf4*, and *irf1_isoform2_*) in SHK-1 cells after IFNa treatment and infection. The panels represent the following conditions: (1) crfb1a knockout (KO) cells infected with ISAV (top panel), (2) crfb1a KO cells infected with IPNV (second panel), (3) crfb5a KO cells infected with ISAV (third panel), and (4) crfb5a KO cells infected with IPNV (bottom panel). Data are shown for the control group (green, non-KO without IFNa), no-KO_IFNa group (blue, non-KO with IFNa), and KO_IFNa group (orange, KO with IFNa). Statistical significance is indicated as *P < 0.05, **P < 0.01, ***P < 0.001. Error bars represent standard deviation (SD). Biological replicates: n=4 for control, n=3 for KO groups.

### Differential gene expression in effector molecules

Effector molecules, like ISGs, are vital for antiviral defense, which mediate immune responses to block viral replication and spread.. Two important ISGs, *isg15* and *Mx*, exhibited minimal changes in all the KO groups after ISAV infection (Figure 5) except significant upregulation of *isg15* in the combined KO group (Figure S F3). However, in IPNV infected groups, both *isg15* and *Mx* showed increasing expression trends in all three KO groups (Figure 5).

**Figure 5:**
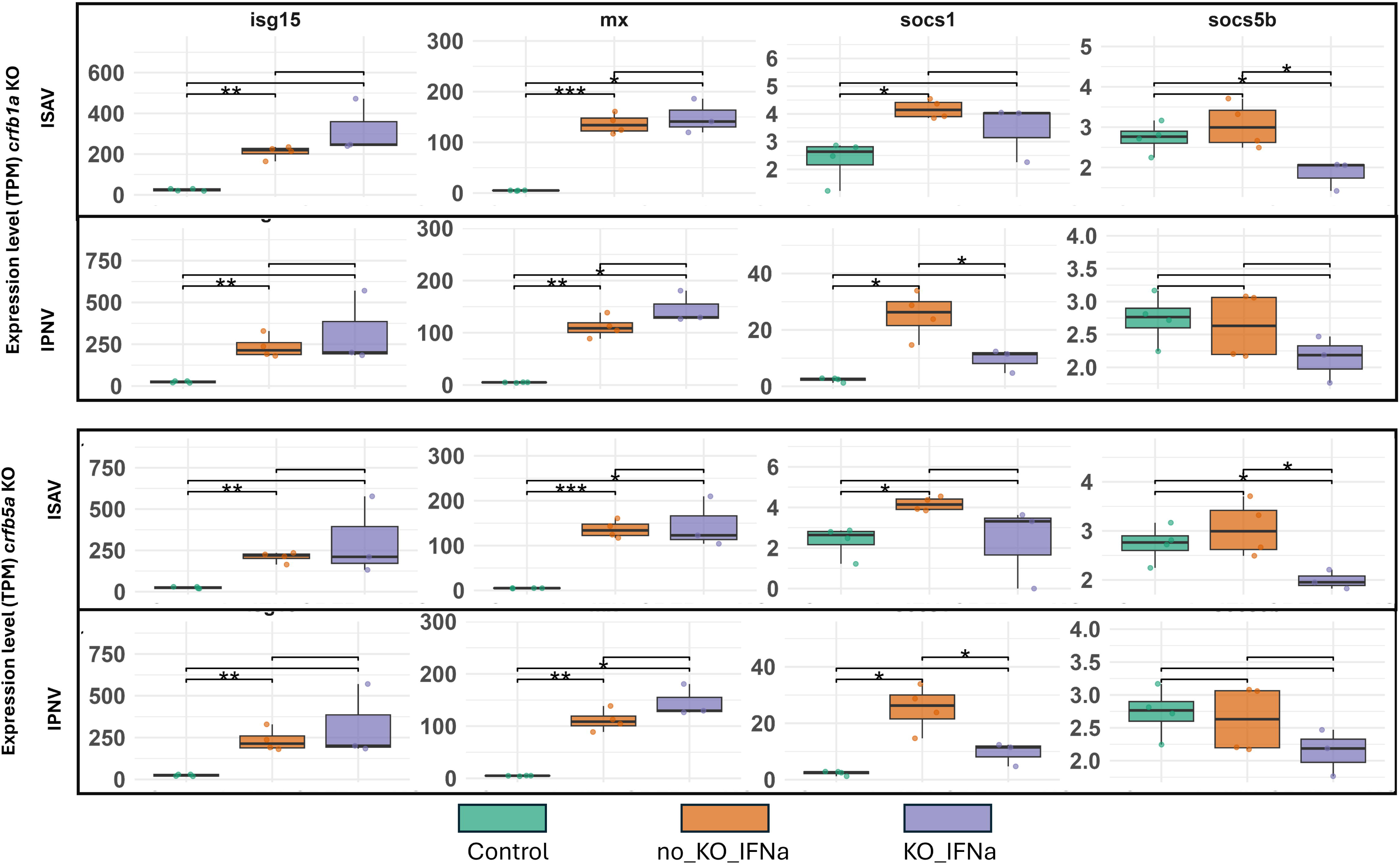
Expression levels (TPM) of of effector molecules (*isg15, Mx, socs1 and socs5b*) in SHK-1 cells after IFNa treatment and infection. The panels represent the following conditions: (1) crfb1a knockout (KO) cells infected with ISAV (top panel), (2) crfb1a KO cells infected with IPNV (second panel), (3) crfb5a KO cells infected with ISAV (third panel), and (4) crfb5a KO cells infected with IPNV (bottom panel). Data are shown for the control group (green, non-KO without IFNa), no-KO_IFNa group (blue, non-KO with IFNa), and KO_IFNa group (orange, KO with IFNa). Statistical significance is indicated as *P < 0.05, **P < 0.01, ***P < 0.001. Error bars represent standard deviation (SD). Biological replicates: n=4 for control, n=3 for KO groups.

In contrast, negative regulator genes like *socs1* and *socs5b* were generally downregulated in all KO groups following ISAV or IPNV infections, with one exception: in the combined KO group after ISAV infection, *socs1* was upregulated. Specifically, in ISAV-infected cells, *socs5b* was significantly downregulated in the *crfb1a* (*p*=0.02) and *crfb5a* (*p*=0.03) KO groups.

Meanwhile, in IPNV-infected cells, *socs1* was significantly downregulated in both the *crfb1a* (*p*=0.04) and *crfb5a* (*p*=0.02) KO groups.

### Pathway Analysis

To understand the cellular response to receptor knockouts, KEGG pathway analysis was performed on DEGs from each KO group (Figure 6). Common DEGs across the three knockout groups are highlighted within the boxed area of the figure. Pathways related to cell motility (regulation of actin), cellular community (focal adhesion), and transport and catabolism (phagosome) were upregulated in both ISAV- and IPNV-infected groups. Conversely, pathways related to signaling pathways (cytokine-cytokine receptor interaction, ECM-receptor interaction) were downregulated in both viral groups.

**Figure 6:**
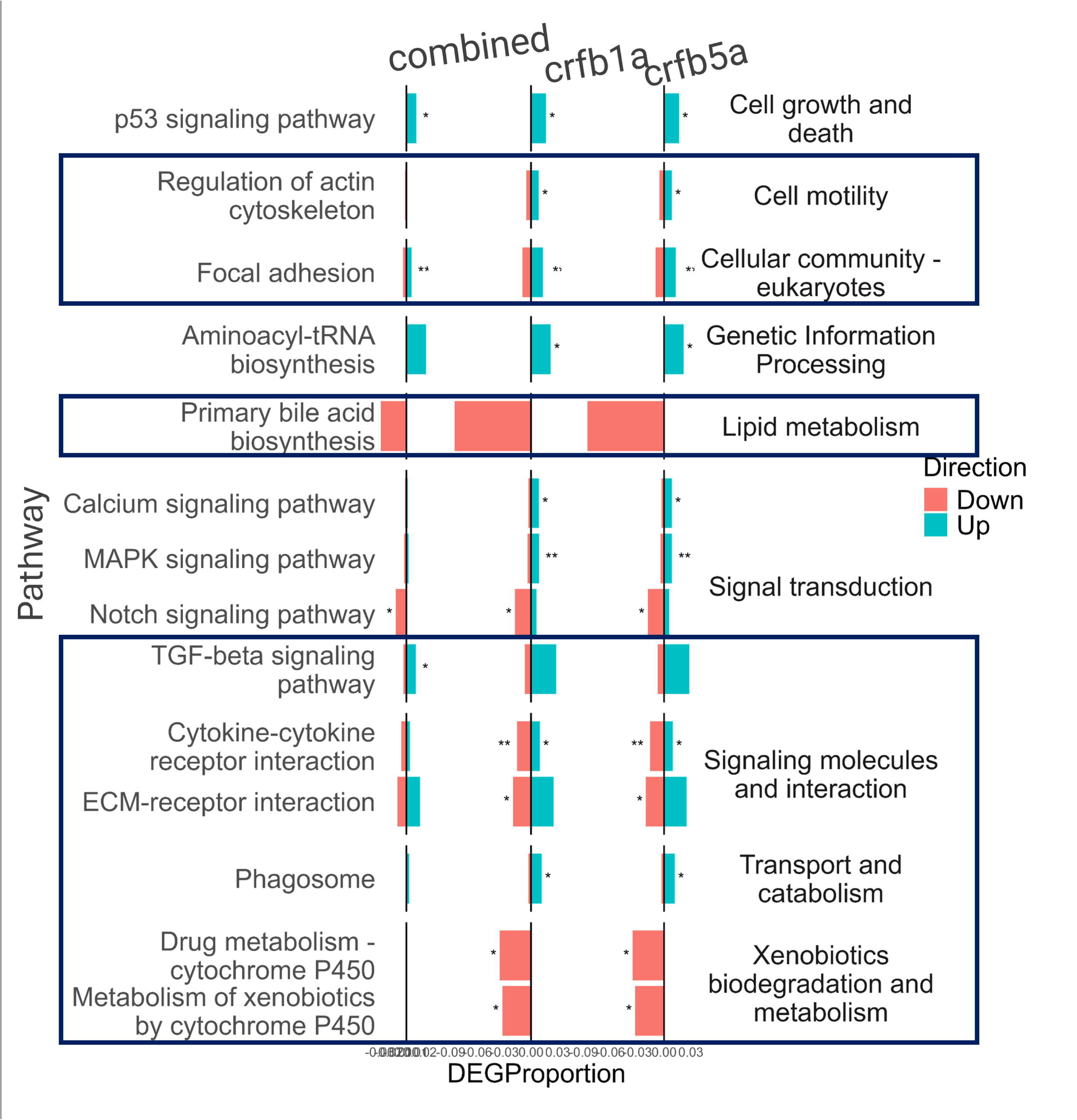
Pathway enrichment analysis illustrating the impact of crfb1a, crfb5a, and combined knockouts on SHK-1 cells infected with Infectious Pancreatic Necrosis Virus (IPNV). Pathways highlighted within the box indicate enrichment in cells infected with both Infectious Salmon Anemia Virus (ISAV) and IPNV. Each bar represents the proportion of differentially expressed genes (DEGs) that are upregulated (red) or downregulated (blue) within various biological pathways. The biological processes are grouped by functional categories. Statistical significance of pathway enrichment is denoted by asterisks (*p < 0.05, **p < 0.01, ***p < 0.001)

## Discussion

The findings presented herein is an important step towards deciphering the different interferon receptors and their mediated antiviral responses. Despite of the similarities between IFN responses in higher and lower vertebrates, the detailed understanding of the signaling mechanism is lacking in non-mammalian vertebrates and much of the available knowledge are generated by extrapolation. Hence, the need for studies addressing the IFN signaling mechanism is eminent. Generating knockout receptors using different methods including CRISPR is a useful approach in this regard. In this study, four candidate IFN receptors were knocked out from SHK-1 cells using CRISPR-Cas9. The selected receptors were the IFNa receptors *crfb1a* and *crfb5a* (Sun et al., 2014) and *ifngr2a*, *il10r2* which interacts with IFN gamma in mammals (Kotenko et al., 2003; Pestka et al., 2004). Along with these single KOs, the candidate IFN receptors were also combinedly KO from SHK-1 together with a fifth candidate receptor (*il10r2)*. Our hypothesis was, complete KO of essential receptors will significantly reduce the genes expression in downstream pathway, signaling cascades and ISGs production even we induce IFNa. Indeed, we observed a downregulation of genes involved the immediate (STAT1, 2 and 6) and downstream (IRF3 and 7) signaling pathway while the effect on the effectors was limited to the SOCS. The effect of the different knockout was however variable. This may be due to the complicated study design which included infection with IPNV and ISAV, made to build up on the findings of our previous study (Reza et al., in review). A simpler design with no infection would have probably resulted in a clearer effect and should be adopted in future studies. Moreover, the different editing efficiencies may have also impacted the results. Hence optimization of the editing protocol to achieve 100% editing is recommended. One approach to do that will be KO by lentiviral transduction and using antibiotic selection, or using Cas9 or guides coupled to GFP and fluorescent cell sorting (Strømsnes et al., 2022).

Our results also highlight difference in IFN receptor mediated responses between IPNV and ISAV which can be helpful in understanding the pathogenic mechanisms and the interplay between the IFN responses and these two viruses. In Atlantic salmon, it was previously shown that infection with both ISAV and IPNV do not significantly induces ISGs, since both viruses have the antagonizing properties (Dahle & Jørgensen, 2019). Studies showed that, ISAV has antagonist protein in segment 7 and s8ORF2 which act as a RNA-silencing suppressor and reduce IFN and ISGs production significantly (García-Rosado et al., 2008; Li et al., 2016; McBeath et al., 2006; Thukral et al., 2018). Similarly, by studying the capability to induce the activation of the Atlantic salmon IFNa1 promoter, it was shown that VP2, VP3, VP4 and VP5 of IPNV has antagonizing properties against the IFNa promoter and were therefore suggested as antagonizing proteins that inhibits the IFNa induction (Lauksund et al., 2015). This gives ISAV and IPNV chance to replicate inside the salmon cells. However, creating an antiviral state can lead to the production of ISGs and give some protection or significantly lower the infection (Jensen & Robertsen, 2002; Xu et al., 2010). Among the different IFNs, induction of recombinant IFNa can significantly induce ISGs and provide protection in salmon cells (Xu et al., 2010). Our previous study (Reza et al., In review) demonstrated that without recombinant IFNa induction, ISAV and IPNV infections did not significantly induce *isg15* or *Mx* expression in SHK-1 cells. However, when treated with recombinant IFNa, the highest expression of these genes was observed two hours post-infection. Building upon these findings, we optimized the experimental conditions, including pre-treatment with IFNa and focusing on the two-hour time point, to effectively study the role of CRISPR-mediated knockouts of IFNa receptors in this study. These optimized conditions allowed for a more precise evaluation of the antiviral effects and pathways influenced by these receptors. In concordance with this, IFN receptor KO did not affect expression of either *Mx* or *isg15* which may be due to the mechanism employed by these two viruses to downplay these responses as previously indicated (Lauksund et al., 2015; Li et al., 2016).

The immediate downstream components of the JAK-STAT pathway, including *stat1b, stat2,* and *stat6*, were significantly downregulated in *crfb1a* KO cells during ISAV infection. *Crfb5a* KO showed a more selective effect, with significant downregulation observed only for *stat2*. In IPNV-infected cells, *stat2* exhibited a downregulation trend in both *crfb1a* and *crfb5a* KO groups. These results demonstrate the involvement of the *crfb1a* and *crfb5a* receptors binding with the IFNa in Atlantic salmon cells, with the knockout of *crfb1a* leading to more pronounced effect compared to *crfb5a*. This aligns with the findings of Sun et al. (2014) who demonstrated that *Mx* promoters were activated in CHSE cells transfected with CRFB1a and CRFB5a, highlighting their involvement in type I IFN receptor-mediated responses (Sun et al., 2014). Additionally, the downregulation of JAK-STAT genes confirms their involvement in the type I IFN signaling pathway, aligning with the canonical model of IFN signaling observed in fish, mammals, and other species (Liu et al., 2024; Majoros et al., 2017; Skjesol et al., 2010). Moreover, upstream components such as *jak1* and *tyk2* remained unaffected which indicate that the receptor KO predominantly impacts downstream signaling and responses.

SOCS genes, known as negative regulators of cytokine signaling, were also notably impacted. In ISAV-infected cells, *socs5b* was significantly downregulated in *crfb1a* and *crfb5a* KO groups, while *socs1* was significantly downregulated in IPNV-infected cells under the same conditions. This aligns with the notion that SOCS gene expression is closely tied to the activity of the JAK-STAT pathway, which was suppressed following IFNa receptor KO. The observed downregulation of SOCS genes suggests that their regulatory function was minimized, likely due to the reduced activation of the JAK-STAT pathway. In addition, the findings reported here support the conserved role of *socs1* and *socs5b* as a key negative regulators of IFN signaling, inhibiting the JAK-STAT pathway and its downstream signaling (Guo et al., 2019; Linossi et al., 2013; Pan et al., 2023).

The results presented also demonstrated distinct patterns of gene regulation between ISAV and IPNV infections. ISAV infection led to pronounced downregulation of multiple STAT family members, accompanied by significant suppression of *socs5b*. In contrast, IPNV infection showed less effects on STAT expression but resulted in significant downregulation of *socs1*. Moreover, *irf1* expression was significantly upregulated only after IPNV infection and downregulated in *crfb1a* and *crfb5a* KO compared to no-KO only after IPNV infection. IRF1 is a key regulatory factors and upregulation of IRF1 was observed in different virus infection, or upon IFNa induction (Feng et al., 2021; Kanazawa et al., 2004). A study in CHSE-214 cell showed that IRF1 expression was upregulated after IPNV infection and further induced the phosphatidylserine receptor (PSR) which help in clearing the apoptotic cells (Kung et al., 2014). IRF1 is also involved in different signaling such as toll like receptor signaling, IFN mediated signaling, IFN-dependent inflammation to increase the host defense against pathogens (Feng et al., 2021). Through additional knockout studies, the role of the abovementioned and other signaling molecules and transcription factors during different virus infections, which was beyond the goal of this study, can be addressed.

Our study showed upregulation of *irf7* in *crfb1a* and *crf5a* KO group after both ISAV and IPNV infection. Also, genes in the effector molecules such as *isg15* and *Mx* showed similar or even higher expression than the no_KO group. ISGs can be induced upon viral infection independently from IFN signaling pathways in different ways (Swaraj & Tripathi, 2024; Tandel et al., 2022). A study explicitly demonstrated that GS2 cells (STAT2 knockout) retained resistance to viral infections despite the absence of the canonical type I IFN-induced ISG response and suggested that alternative pathways contributed to antiviral defense (Dehler et al., 2019). During induction of type I IFN response, IRF7 acts as a master regulator (Honda et al., 2005; Ma et al., 2023). Further, it was shown that IRF7 is capable of directly binding with the ISREs of some ISGs and induce it in JAK-STAT independent pathways (Schmid et al., 2010). Hence, it is possible that, despite the downregulation of JAK-STAT IRF7 has been induced in JAK-STAT independent pathways in response to virus infections.

The KEGG pathway analysis highlights significant cellular responses and upregulation of actin regulation, focal adhesion, phagosome reflects an enhanced phagocytic activity (Jaumouillé & Waterman, 2020; May & Machesky, 2001). This enhancement likely serves to promote the efficient clearance of viral particles and cellular debris, thereby compensating for compromised immune signaling pathways resulting from receptor knockouts.

In our study, CRISPR-Cas9 KO were done by using ribonucleoprotein (RNP) complex to create the KOs. This is an established method to create efficient KO in Atlantic salmon cells (Pavelin et al., 2021). The efficiency of different KO varied and was not 100%, which indicates that the effect may not fully reflect the impact of a complete knockout. The study showed clear separation of *crfb1a*, *crfb5a*, combined KO group from the IFN induce group suggest their response after KO. However, for IFNGR2a that was not case. Since this is a receptor in IFN gamma, in fish also it may not be triggered after IFNa induction.

There are some factors we need to consider while explaining the results. Because of the whole genome duplication in fish and an additional duplication in salmonoid there can be four potential paralogous genes for each genes (Lien et al., 2016; Pasquier et al., 2016). We have selected one for each based on the Sun et al., (2011) study. However, the paralogous gene may act as compensatory gene in the absence of one or more of other paralogues. Moreover, for *crfb5a*, gRNA was designed in the 4^th^ exon of the gene. There were two more alternative TSS after the designed gRNA position which may also be transcribed and could potentially produce functional protein. Hence this study underscores the complexity of IFN signaling in Atlantic salmon and the possible compensatory mechanisms that can be exploited after editing. Future research should investigate paralogous gene functions and alternative pathways to enhance disease resistance strategies in aquaculture.

## Author contributions

All authors contributed to the study conception and design. Material preparation, data collection and analysis were performed by Mohammad Ali Noman Reza. The first draft of the manuscript was written by Mohammad Ali Noman Reza and all authors commented on previous versions of the manuscript. All authors read and approved the final manuscript.

