## Supplementary file for "Investigating antiviral pathways in Atlantic salmon cells through interferon receptor knockouts via CRISPR-Cas9"

| gRNA Name | gRNA Sequence | After 7 days  Indel % | After 14 days  Indel % | After 21 days  Indel % | After 31 days  Indel % | After 92 days  Indel % | After 104 days  Indel % |
| --- | --- | --- | --- | --- | --- | --- | --- |
| crfb1a | GAAGAGGGCAGGAGAGACAA | 65 | 62 | 61 | na | 51 | na |
| crfb5a | GAAGTTACAGTGGGTATGTA | 78 | 88 | 85 | 83 | na | 86 |
| ifngr2a | GCTGCAGGAATCAGCTGGGT | 90 | 76 | 84 | 90 | na | 90 |
| il10rb | TCACCTGTACTCTGCTCTGT | 95 | 97 | 97 | na | 96 | na |

### S. T1 Knockout Efficiency of Individual gRNAs Over Time

### S. T2 Knockout Efficiency of Combined IFN Receptor Genes Over Time

| Date | crfb1a Indel % | crfb5a Indel % | i10r2  Indel % | Ifng2a  Indel % | I10rb  Indel % |
| --- | --- | --- | --- | --- | --- |
| After 7 days | 52 | 64 | 63 | 97 | 96 |
| After 31 days | 52 | 86 | na | 90 | 96 |
| After 38 days | na | 66 | na | 98 | 96 |

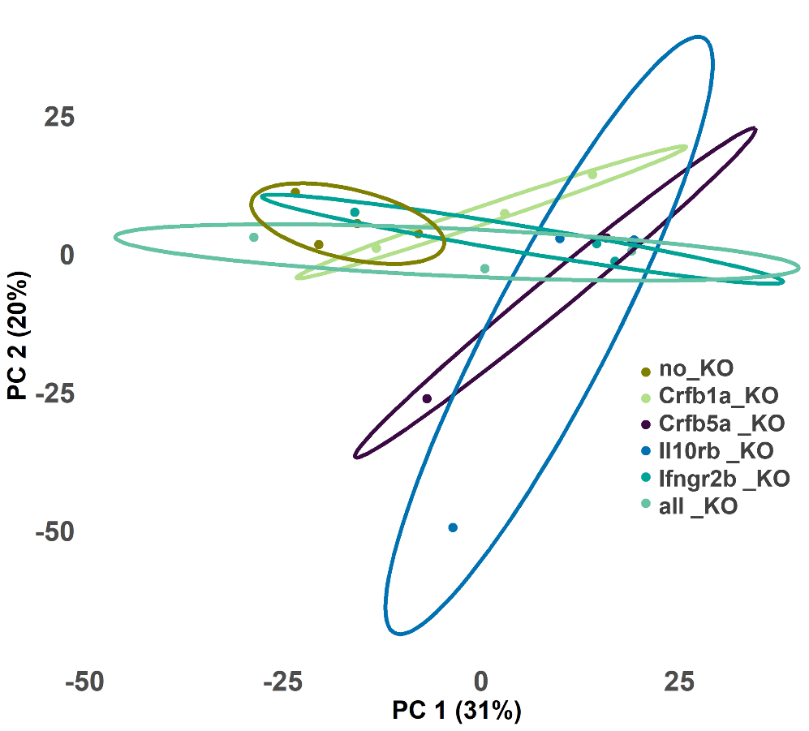

S F1. RNA Sequencing and transcriptomic analysis of SHK-1 Cells KO after IFNa receptors under ISAV and IPNV infections. Principal component analysis (PCA) displaying gene expression clustering among SHK-1 cells in ISAV and IPNV cells with individual KO of and combined IFN receptor genes post-infection. PC1 and PC2 represent the indicated percentage of the variance, respectively.

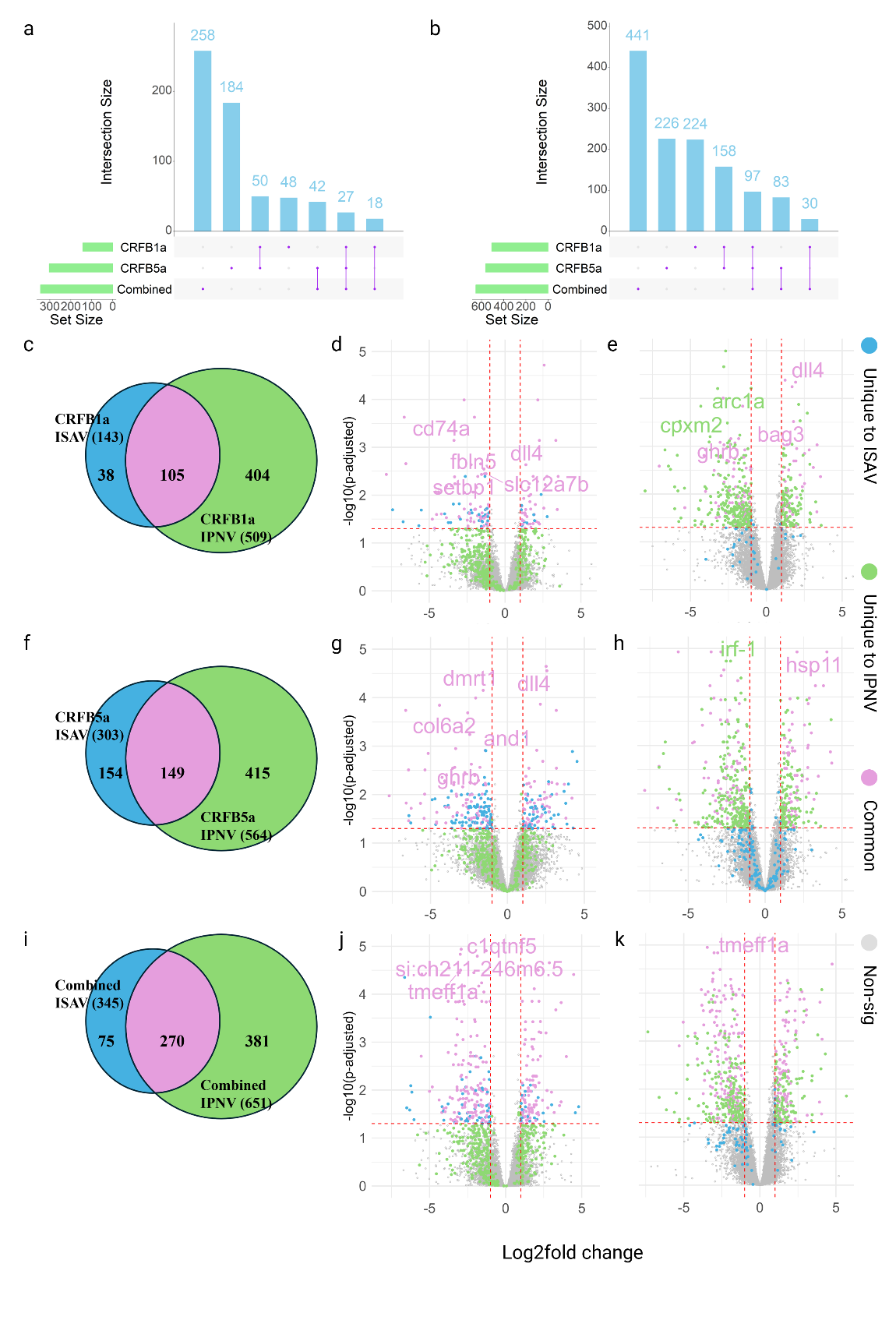

S F2. Upset plots, Venn diagrams, and volcano plots illustrating the differential expression of genes (DEGs) in SHK-1 cells knocked out for CRFB1a, CRFB5a, and combined IFN receptor genes post-infection with ISAV and IPNV.

a-b. Upset plots show the intersection of DEGs identified in SHK-1 cells with knockouts for CRFB1a, CRFB5a, and the combined knockout following ISAV infection (a) and IPNV infection (b). Bars represent the intersection size of DEGs across different knockout conditions, while the horizontal bars at the bottom denote the total set size for each condition.

c. Venn diagram displaying the overlap of DEGs between ISAV and IPNV infections in SHK-1 cells with CRFB1a knockout.

d-e. Volcano plots showing unique and common DEGs in the CRFB1a knockout with ISAV infection (d) and the CRFB1a knockout with IPNV infection (e).

f. Venn diagram representing the overlap of DEGs between ISAV and IPNV infections in the CRFB5a knockout group.

g-h. Volcano plots for the CRFB5a knockout group, displaying DEGs following ISAV infection (g) and IPNV infection (h).

i. Venn diagram showing the overlap of DEGs between ISAV and IPNV infections in the combined knockout group.

j-k. Volcano plots for the combined knockout group, illustrating DEGs following ISAV infection (j) and IPNV infection (k).

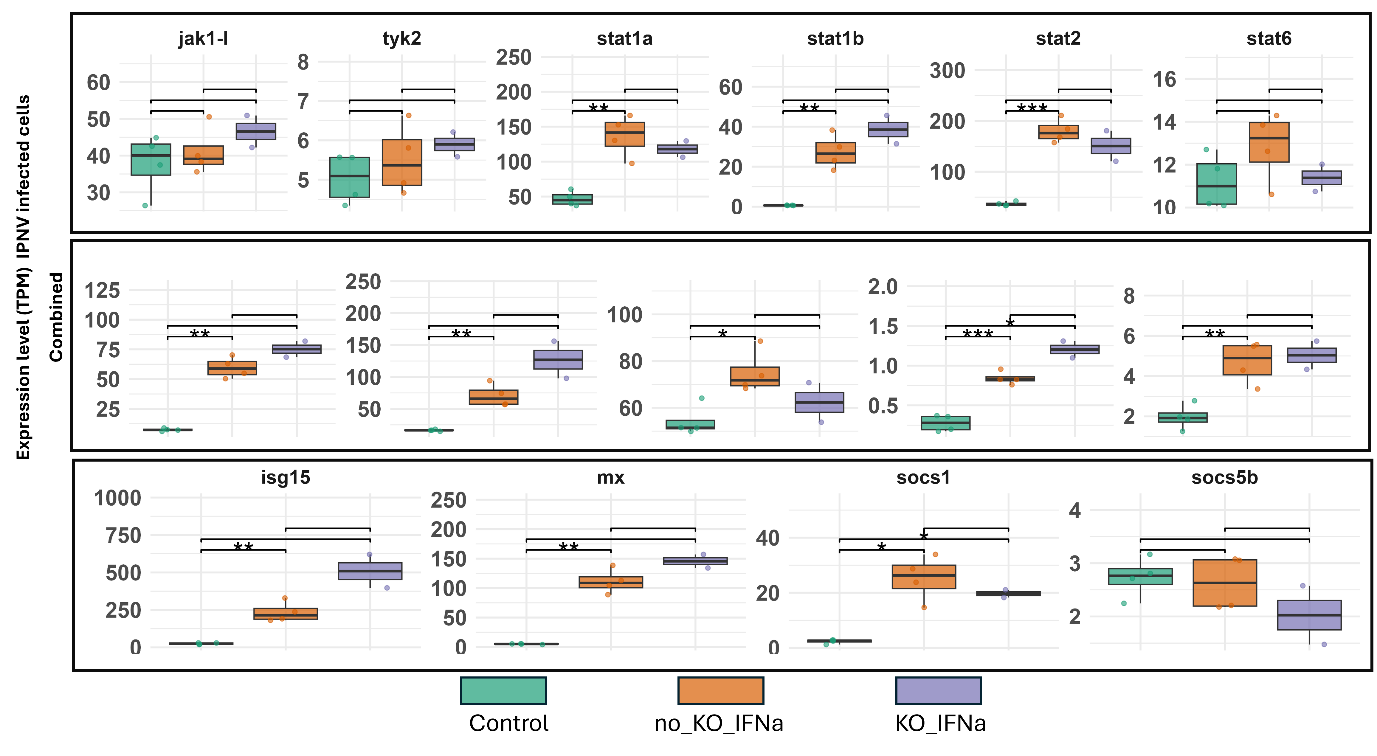

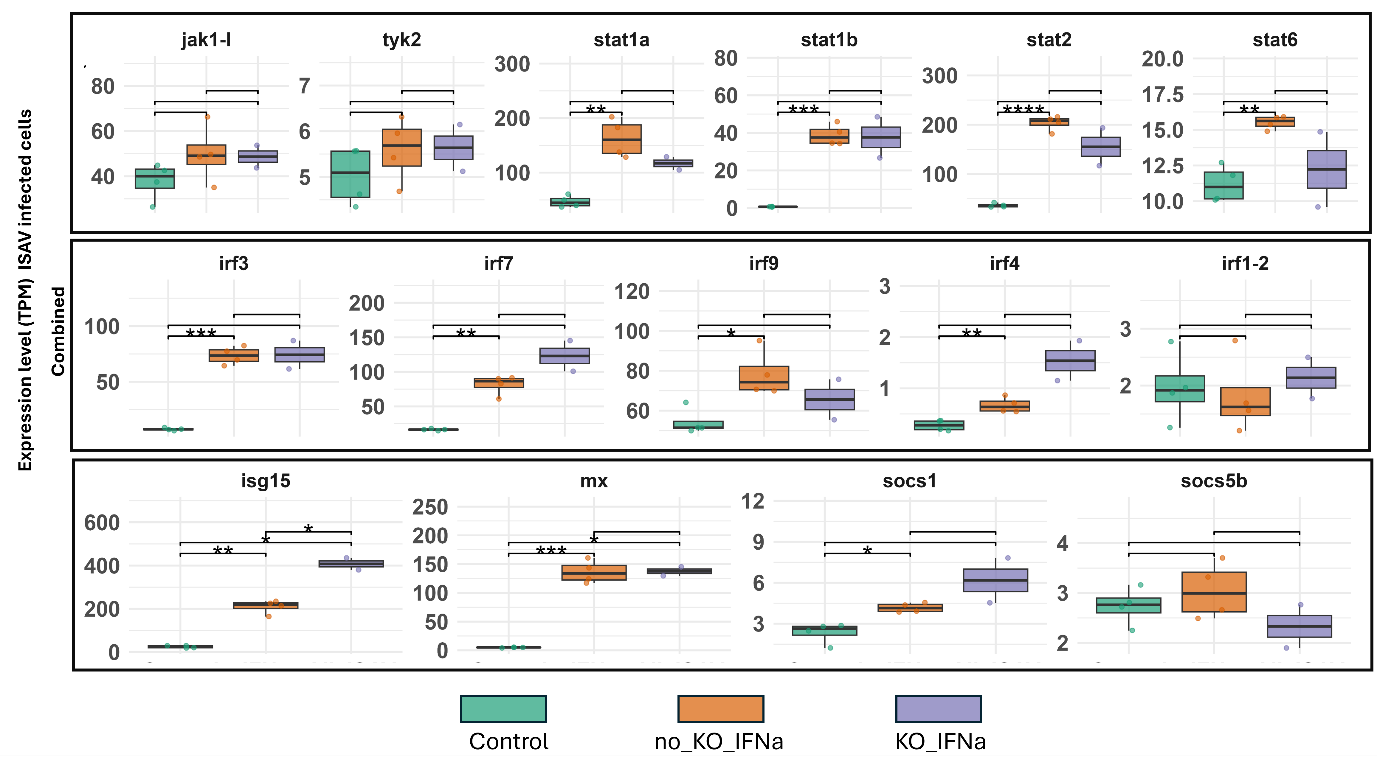
Figure S F3: Expression levels (TPM) of key genes (jak1-l, tyk2, stat1a, stat1b, stat2, stat6, isg15, mx, socs1, and socs5b) in SHK-1 cells infected with ISAV and treated with IFNa. Data represent the combined receptors knockout (KO) group for *crfb1a*, *crfb5a*, *il10rb*, *ifngr2a*, *il10r2*. The groups shown are control (green, non-KO without IFNa), no_KO_IFNa (blue, non-KO with IFNa), and KO_IFNa (orange, combined KO with IFNa). Statistical significance is indicated as *P < 0.05, **P < 0.01, ***P < 0.001. Error bars represent standard deviation (SD). Biological replicates: n=4 for control, n=3 for KO groups.

Figure S F4: Expression levels (TPM) of key genes (jak1-l, tyk2, stat1a, stat1b, stat2, stat6, isg15, mx, socs1, and socs5b) in SHK-1 cells infected with IPNV and treated with IFNa. Data represent the combined receptors knockout (KO) group for *crfb1a*, *crfb5a*, *il10rb*, *ifngr2a*, *il10r2*. The groups shown are control (green, non-KO without IFNa), no_KO_IFNa (blue, non-KO with IFNa), and KO_IFNa (orange, combined KO with IFNa). Statistical significance is indicated as *P < 0.05, **P < 0.01, ***P < 0.001. Error bars represent standard deviation (SD). Biological replicates: n=4 for control, n=3 for KO groups.
